# ACME: Pan-specific peptide-MHC class I binding prediction through attention-based deep neural networks

**DOI:** 10.1101/468363

**Authors:** Yan Hu, Ziqiang Wang, Hailin Hu, Fangping Wan, Lin Chen, Yuanpeng Xiong, Xiaoxia Wang, Dan Zhao, Weiren Huang, Jianyang Zeng

## Abstract

Prediction of peptide binding to MHC molecules plays a vital role in the development of therapeutic vaccines for the treatment of cancer. Although numerous computational methods have been developed to this end, several challenges still remain in predicting peptide-MHC interactions. Many previous methods are allele-specific, training separate models for individual alleles and are thus unable to yield accurate predictions for those alleles with limited training data. Despite that there exist several pan-specific algorithms that train a common model for different alleles, they only adopt simple model structures that generally have limited performance in capturing the complex underlying patterns of peptide-MHC interactions. Here we present ACME (Attention-based Convolutional neural networks for MHC Epitope binding prediction), a new pan-specific algorithm to accurately predict the binding affinities between peptides and MHC class I molecules, even for those new alleles that are not seen in the training data. Extensive tests have demonstrated that ACME can significantly outperform other state-of-the-art prediction methods with an increase of the Pearson Correlation Coefficient by up to 23 percent. In addition, its ability to identify strong-binding peptides has been experimentally validated. Moreover, by integrating the convolutional neural network with attention mechanism, ACME is able to extract interpretable patterns that can provide useful and detailed insights into the binding preferences between peptides and their MHC partners. All these results have demonstrated that ACME can provide a powerful and practically useful tool for the studies of peptide-MHC class I interactions.

## 1 Introduction

Presentation of antigen peptides by major histocompatibility complex (MHC) class I molecules plays a vital role in initiating an immune response to identify and kill cancer cells. Peptides derived from mutant proteins in cancer cells can be engineered into cancer vaccines, which can stimulate a specific immune response against the cancer cells presenting these antigen peptides [1–3]. However, in most cases, for a peptide to effectively elicit an immune response, it needs to bind to an MHC molecule with a sufficiently high affinity (IC_50_*<*500 nM), making peptide-MHC binding the most crucial and selective step in antigen presentation [4]. Hence, identifying the peptides that can be presented by MHC molecules is a vital step in developing effective therapeutic cancer vaccines [5]. This underscores the importance of developing *in silico* algorithms that can accurately predict the binding affinities of peptides to MHC molecules.

In the past two decades, numerous computational methods have been proposed for this purpose [6–15]. Two general strategies (i.e., allele-specific and pan-specific predictors) have typically been adopted by the computational methods presented so far. (1) The allele-specific predictors generally train one model for each MHC allele. For example, NetMHC, one of the most widely used allele-specific models, employs a conventional feed-forward neural network with a single hidden layer to predict peptide-MHC interactions for individual alleles [11, 12]. When training these allele-specific methods, we have to split the binding data into individual smaller datasets, each specific for a single allele, making it difficult to derive accurate predictions for those data-deficient alleles. (2) On the other hand, the pan-specific methods take both peptide and MHC sequences as input features, thus allowing to pool all the data for different alleles together and train one common model for all alleles. For instance, the widely used method, NetMHCpan, uses a simple neural network model with one hidden layer and takes information extracted from both peptide and MHC sequences as its input to perform the pan-specific prediction task [6–9]. However, such neural networks with a single hidden layer are generally not sophisticated enough to capture all the intrinsically complex patterns in the data. The recently proposed method, ConvMHC, uses deep convolutional networks [14] to address this problem. Unfortunately, it cannot integrate the binding data of peptides with different lengths, and is only applicable to 9-mer peptides.

Despite the progress made so far, there is still a need for developing a more precise method for peptide-MHC binding prediction, because weak binders greatly outnumber strong ones in this scenario, leading to a considerable number of false positives when identifying antigen peptides. In addition, the prediction results from most of the previous methods lack interpretability. Although there existed several studies on the interpretable binding patterns between peptides and MHC modules [16–18], they only focused on a limited number of alleles. Thus, it becomes increasingly necessary to develop a model that can not only make accurate binding prediction but also reveal interpretable patterns to characterize peptide-MHC interactions. Recently, the attention mechanism has been used in natural language processing [19, 20], computer vision [21] and bioinformatics [22] to produce interpretable results. It assigns different weights to individual positions of the input, so that the model can focus on the most crucial features to perform better prediction. In sequence analysis, the attention values (weights) for individual sites (e.g., base pairs or amino acid residues) derived by the embedded attention mechanism allow one to spot those important sites that significantly contribute to the final predictions. Hence, here attention is a suitable technique for interpreting the binding patterns of MHC molecules.

In this article, we propose a new deep learning based method, called ACME (Attention-based Convolutional neural networks for MHC Epitope binding prediction), for the pan-specific prediction of the binding affinities between peptides and MHC class I molecules. We combine a deep convolutional neural network with an attention module to build an accurate and interpretable prediction model, in which features of different levels are extracted from multiple layers of the convolutional network, and then integrated together to effectively capture the intrinsic characteristics of peptide-MHC binding. Extensive tests on existing benchmark and external datasets have demonstrated that ACME can significantly outperform other existing state-of-the-art prediction methods with an increase of the Pearson Correlation Coefficient by up to 23 percent. More importantly, we have also shown that ACME can be effectively used to make accurate predictions for those new MHC alleles that are not seen in the training data. In addition, the ability of ACME to identify strong binding peptides has been well validated experimentally. Moreover, comprehensive analyses have indicated that ACME can extract interpretable molecular interaction patterns and generate accurate peptide binding motifs for different MHC alleles, including those MHC alleles with scarce or no available experimental data. Overall, our model can provide a powerful and useful tool for predicting peptide-MHC binding as well as investigating the underlying patterns of this biological process. ACME is available as an open source software at https://github.com/HYsxe/ACME

## 2 Materials and methods

### 2.1 Datasets

The widely used IEDB MHC class I binding affinity dataset was downloaded from the IEDB website (http://tools.immuneepitope.org/mhci/download/) [23, 24]. We first carried out a five-fold cross-validation procedure on this dataset to evaluate the performance of our model and other state-of-the-art prediction methods. Afterwards, we combined this IEDB dataset with another dataset obtained from Pearson et al. [25], to train a final version of our model (Redundant data were removed). To test if our final model can be generalized to other datasets, we further tested our model on the data obtained from the IEDB weekly benchmarking website (http://tools.iedb.org/auto_bench/mhci/weekly/) [26], which had also been previously used to test other methods as well [14, 15]. In particular, the datasets published within the last three years were chosen. These datasets were not used in model optimization and training of our final model. The sequences of different MHC molecules were obtained from the IPD-IMGT/HLA Database (ftp://ftp.ebi.ac.uk/pub/databases/ipd/imgt/hla/,hla_prot.fasta) [27]. More details about dataset preprocessing and filtering can be found in Supplementary Notes Section 1.1.

### 2.2 Data encoding

The input data to our prediction framework ACME include peptide and MHC sequences, which are both encoded based on the standard BLOSUM50 scoring matrix [28], with each residue encoded by the corresponding row of this matrix. In our framework, the elements of the BLOSUM matrix are also divided by a scaling factor to facilitate model training, which was set to 10 after hyperparameter tuning using a grid search scheme. Each peptide is encoded into a 24*×*20 matrix in a pair-end manner (Figure 1a). In particular, the first 12 rows of the input matrix encode the input peptide with the N terminus aligned to the first row, while the last 12 rows encode the peptides with the C terminus aligned to the last row. For those peptides shorter than 12 residues, the vacancies in the middle are filled with zero padding. For instance, a peptide sequence of length 9 is encoded into the 1^*st*^-9^*th*^ and 16^*th*^-24^*th*^ rows, in which the 1^*st*^ and 16^*th*^ row encode the N terminus while the 9^*th*^ and 24^*th*^ rows encode the C terminus. In this case, the 10^*th*^ to 15^*th*^ rows are all padded with zeros.

**Figure 1:**
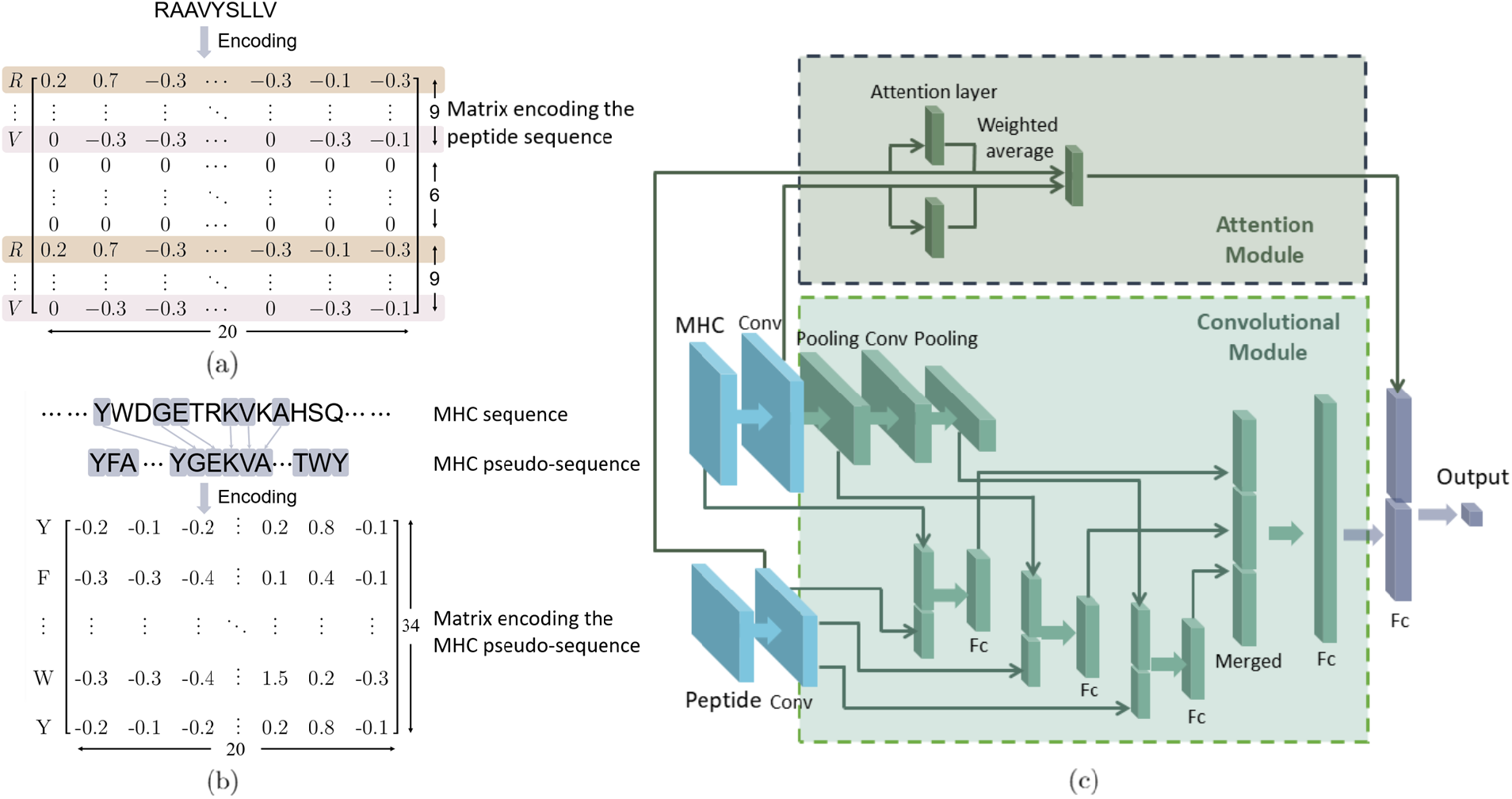
Schematic illustration of the ACME framework. a Our scheme for encoding peptide sequences. Each peptide sequence is encoded into a 24 *×* 20 matrix in a pair-end manner. b Our scheme for encoding MHC sequences. For each MHC sequence, the 34 residues that are in close spatial proximity with the peptide partner (*<* 4.0 Å) are selected as pseudo-sequence and then encoded into a 34 *×* 20 matrix. c The architecture of the deep neural networks employed in our model. MHC and peptide sequences are encoded and used as input. After initial feature extraction, the encoded feature representations are channeled into both attention and convolutional modules. The output of these two modules are then combined to make the final prediction.

When encoding the input MHC sequences, we only consider those residues that are in close contact with the peptide (i.e., within 4.0 Å distance). These residues form a short sequence, called the MHC pseudo-sequence, as defined in [6]. In particular, 34 such residues are chosen and encoded into a 34 *×* 20 matrix, where each residue is represented by the corresponding row in the BLOSUM50 matrix (Figure 1b).

The peptide-MHC binding affinities, which were represented by the IC_50_ values in nM units, were first transformed using the equation 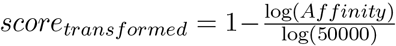, as previously described in [29], and then used as the label values to train and test our model.

### 2.3 Network architecture

Our model ACME mainly consists of two modules, including a convolutional module and an attention module (Figure 1c). ACME first passes the encoded peptide and MHC pseudo-sequences through one convolutional layer for initial feature extraction, and then sends the extracted feature maps into both convolutional module and attention modules. After the that, the outputs of these two modules are combined together to make the final prediction. The model is implemented using Keras [30]. Below we will describe each part of the model in detail.

#### 2.3.1 Initial feature extraction

The encoded peptide and MHC feature representations as input to our model are first processed separately by a one-dimensional convolutional layer with rectified linear unit (ReLU) activation [31] for initial feature extraction. The convolutioal layers mentioned hereafter all use ReLU activation. The convolutional layer outputs a feature map matrix *F*, in which each column corresponds to one convolutional filter and each row represents a position of the input, that is,

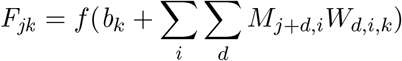

where *M* stands for the input matrix (24 *×* 20 for the peptide sequence or 34 *×* 20 for the MHC pseudo-sequence). *f* stands for the ReLU activation function, i.e., *f*(*X*)_*jk*_ = *max*{*X_jk_*, 0}. *W_d,i,k_* stands for the element in the *d^th^* row and the *i^th^* column of the *k^th^* filter, and *b_k_* stands for the corresponding bias. Note that the ranges of *d* and *j* may vary due to different padding results. The resulting feature maps for the peptide and MHC sequences, extracted by the first convolutional layers, are then separately forwarded into both convolutional and attention modules.

#### 2.3.2 Convolutional module

The convolutional module is mainly designed to extract features of different levels and then integrate them to make prediction. In our convolutional module (Figure 1c), the MHC feature map *F* first goes through a max-pooling layer, followed by another round of one-dimensional convolution and max-pooling operations, while the peptide feature map does not go through further convolution and pooling operations mainly due to its short sequence length (i.e., 9-12 residues). The MHC features extracted after 0, 1 and 2 rounds of convolution and max-pooling operations are then concatenated separately with the peptide feature map after one convolution layer, thus preserving the MHC features of different levels. Here, each concatenated layer is also connected to a separate fully connected layer. The outputs of these three fully connected layers are then concatenated together and forwarded into two consecutive fully connected layers. The output of the latter is the output of the convolutional module. More details on the network architecture can be found in Supplementary Notes Section 1.2.

#### 2.3.3 Attention Module

The attention module in our framework cooperates with the convolutional module to extract interpretable binding patterns (i.e., motifs). In particular, it assigns different weights to the feature vectors corresponding to individual residue positions and then computes their weighted average to facilitate prediction. As a result, it learns to assign higher weights to those residues that are more important in predicting peptide-MHC interactions.

The structure of the attention module is shown in Figure 2. Each row vector *F_j_* in the feature map matrix *F* is taken as the input to the attention layer and generates a real number *w_j_*. In other words, the attention layer maps the feature map matrix *F* into an attention weight vector *w*. Then each element *w_j_* of *w* is converted into 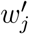 through softmax normalization. Then we use the row vectors to compute their weighted average *F_avg_*, with each 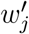 being the weight of the corresponding *F_j_*, that is,

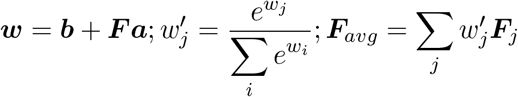

where *a* stands for the weights of the attention layer and *b* is the corresponding bias term. Here, *F_avg_* is the output of the attention module, which will be further combined with the output of the convolutional module to produce the final prediction.

**Figure 2:**
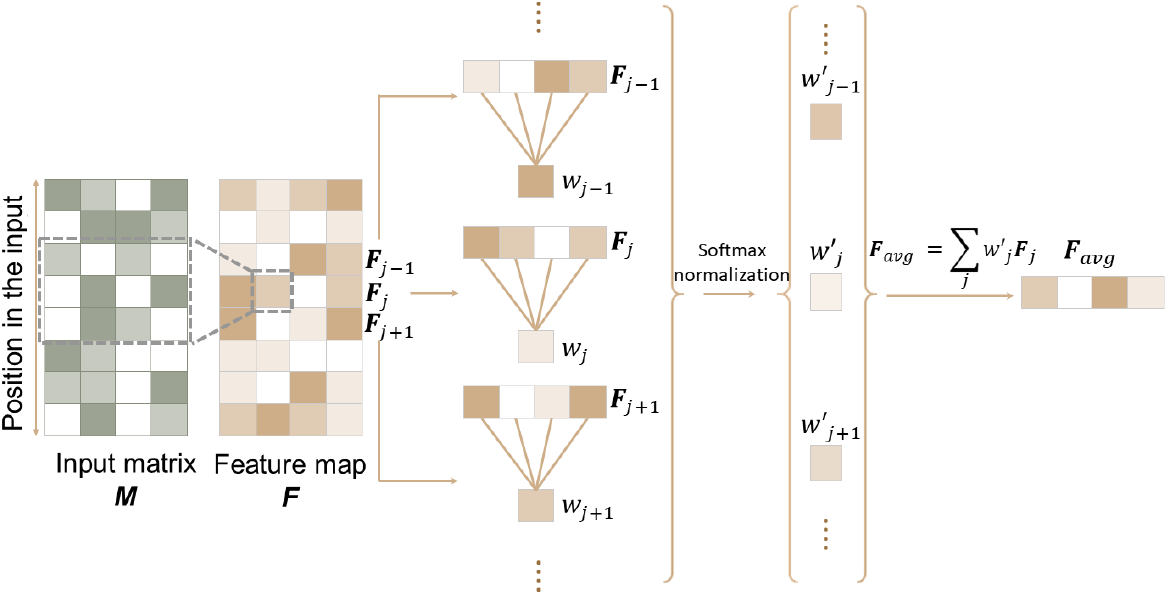
Schematic illustration of the attention module. The convolutional layer extracts a feature map *F* from the input matrix. Each row vector *F_j_* of this feature map *F* corresponds to a position in the input sequence. *F_j_* is used as the input to the attention layer to generate its positional weight *w_j_*, which becomes 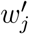 after softmax normalization. *F_avg_* stands for the final weighted average over all row vectors. More details of the attention module can be found in Section 2.3.3.

#### 2.3.4 Performing the final prediction

The outputs of the convolutional module and the attention module are concatenated together and then forwarded into a fully connected layer to perform the final prediction (Figure 1c). It is worth noting that since the outputs of both modules are used to make prediction, the attention module might not be able to account for all the features involved in peptide-MHC binding. Nevertheless, as we will show in the Results section, the attention module can indeed help to capture useful and interpretable patterns that well characterize peptide-MHC binding and provide useful insights into understanding their interactions.

### 2.4 Network training

To determine the hyperparameters of our model, we split the IEDB dataset into a 95% training set and a 5% validation set. The model was optimized using the training set, and its optimal hyperparameters were determined through a grid search approach on the validation set. The performance was evaluated mainly using the Pearson correlation coefficient (PCC), while the Area under the Receiver Operating Characteristics curve (AUROC) was also used as an additional metric to evaluate the prediction results. For the calculation of AUROC, experimentally determined affinities were converted into one or zero labels using a threshold of 500 nM. Each model consists of an ensemble of *n* networks, where *n* = 5 in five-fold cross-validation and *n* = 25 in the final version of our model. The average prediction score over all the networks in the ensemble is used as the final output. We use the mean squared error as the loss function. In addition, the Adam optimizer [32] is used to compute stochastic gradient descent in the training procedure.

After hyperparameter calibration, we ran a five-fold cross-validation procedure on the training set mentioned above to evaluate the predictive performance of our model. Five trials of five-fold cross-validation were conducted, each with a different random initialization and their average result was reported. More details can be found in Supplementary Notes Section 1.3.

### 2.5 Testing the generalizability of ACME

To prove the generalization ability of our model, we also chose the most recent IEDB weekly datasets as external test sets to evaluate our model. Before this test, we augmented the original IEDB training data by also incorporating another dataset from [25] and applied a bootstrapping-like strategy to train a more reliable version of our model. More specifically, the whole combined dataset was randomly divided into five subsets of approximately the same size. Then every four subsets were used to train a separate network, which resulted in five trained models in total. We performed such a process with random partitions of the data five times, and overall we obtained an ensemble of 25 trained networks, whose average prediction result was used as the final output. We then evaluated this final version of our model on 30 datasets of human MHC molecules, which are encoded by human leukocyte antigen type A and B genes (HLA-A and HLA-B). Here we mainly used the Spearman Rank Correlation Coefficient (SRCC) as our performance metric. The performances of five trials were averaged to obtain the final result. We also compared our prediction results to those of previous methods, whose SRCC scores are available on the IEDB weekly benchmarking website (http://tools.immuneepitope.org/auto_bench/mhci/weekly/).

We further tested whether our model can generalize what it has learned to make accurate prediction for those alleles without any training data. Here, for each allele of interest, we removed all the associated binding data from the training set and then used the remaining data to train our model. In particular, data of all other alleles were split into five subsets of approximately equal sizes. We then trained one network on every four subsets, resulting in an ensemble of five networks which takes the average prediction over the five networks as its final output. After that, we used this ensemble to make binding prediction for the allele of interest. Performances were measured in PCC.

### 2.6 Experimental validation

We experimentally validated the ability of ACME to identify neo-antigens derived from somatic mutations in tumors. HLA-A*02:01 and HLA-B*27:05 were chosen as representative MHC alleles with high and low population frequencies, respectively. We selected the most frequent 5000 somatic mutations in human cancer from the COSMIC database [33] and then generated the sequences of all the 9-mer peptides that may potentially present these mutations. For each allele, we used ACME to predict the binding affinities of these peptides and selected the 25 peptides with the highest predicted binding affinities and 25 with the lowest predicted binding affinities for the downstream experimental validation.

We next experimentally measured the actual binding affinities for the above-chosen peptides. For HLA-A*02:01, T2 cells (5 *×* 10^5^ mL^−1^) were incubated with *β*_2_m (3 µg mL^−1^, Sigma) and each peptide (10 µg mL^−1^, Sangon Biotech) at 4 °C for 4 h. The cells were stained with mouse anti-HLAA2 antibody (abcam, ab79523) to measure the expression of HLA-A2 by flow cytometry. The mean fluorescence intensity (MFI) of HLA-A2 was used to indicate the binding affinity of each peptide with HLA-A2 molecules. T2 cells with *β*_2_m only were used as a negative control. T2 cells with *β*_2_m and peptide OVAL235, which had been previously reported to bind to HLA-A2 molecules [34] strongly, were used as a positive control.

For HLA-B*27:05, we used the ProImmune REVEAL MHC-peptide binding assay to determine the binding affinities of the chosen peptides. This assay evaluates the binding affinity of a peptide to an MHC molecule by assessing its ability to stabilize the MHC complex. When the complex is bound and stabilized by the peptide, the MHC protein and *β*_2_m form a native conformation that can be recognized by a labeled antibody and thus generate a positive signal. The percentage of this signal relative to the reference signal generated by a known T cell epitope (positive control) is defined as the ProImmune REVEAL binding score and used to reflect the binding affinity of this peptide.

### 2.7 Generating the binding motifs for different MHC alleles

To better illustrate the underlying biological principles captured by our model, we used ACME to generate binding motifs for different alleles. In particular, we hypothesized that if a certain amino acid type at a specific site contributes significantly to binding, then it should appear more often among strong binders and also be associated with a higher attention score. We use a matrix *A* to represent such an enrichment of attention, where the element in the *i^th^* row and *j^th^* column corresponds to the summed attention score of the *i^th^* amino acid type at the *j^th^* position in the selected peptides, that is,

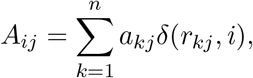

where *n* represents the total number of strong binders and *a_kj_* stands for the attention score assigned to the *j^th^* position of the *k^th^* peptide. Here, *r_kj_* stands for the amino acid type at the *j^th^* position of the *k^th^* peptide. For example, *r_kj_* is 0 for the first type of residue (alanine) and 19 for the last type of residue (valine). Besides, *δ*(*x, y*) is a binary indicator function defined by

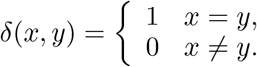

For each allele, we selected the top 1000 peptides with the highest predicted binding affinities (*n* = 1000) from 10,000 random ones and then generated the corresponding matrix A to identify important residues that significantly contribute to MHC-peptide binding. To further investigate the residues with the potential to severely destabilize peptide-MHC complexes, we also selected the bottom 1000 peptides with the lowest predicted binding affinities from 10,000 random ones and also generated their corresponding motifs.

## 3 Results

### 3.1 Five-fold cross-validation on the IEDB MHC class I binding affinity dataset

To evaluate the prediction performance of ACME, we first carried out a five-fold cross-validation on the IEDB dataset for individual alleles and different peptide lengths, including 11-mers, 10-mers and 9-mers (Section 2.4). We compared the performance of our model to that of the widely used state-of-the-art method NetMHCpan [8]. Note that NetMHCpan 3.0 was used here because it was the latest version of NetMHCpan with detailed performance data published for each allele and peptide length.

We mainly used the Pearson Correlation Coefficient (PCC) to assess the predictive abilities of different models. Compared to the previously reported performance of NetMHCpan 3.0, ACME achieved significantly better prediction results, with the average increase of PCC by 11, 5.3 and 3.4 percent for 11-mer, 10-mer and 9-mer, respectively (Figure 3). In addition, we also formulated the peptide-MHC binding problem as a binary classification task and used the Area under the Receiver Operating Characteristic curve (AUROC) to evaluate the prediction performance of ACME. The comparison results based on this metric also demonstrated that ACME significantly outperformed NetMHCpan 3.0 (Supplementary Table 1).

**Figure 3:**
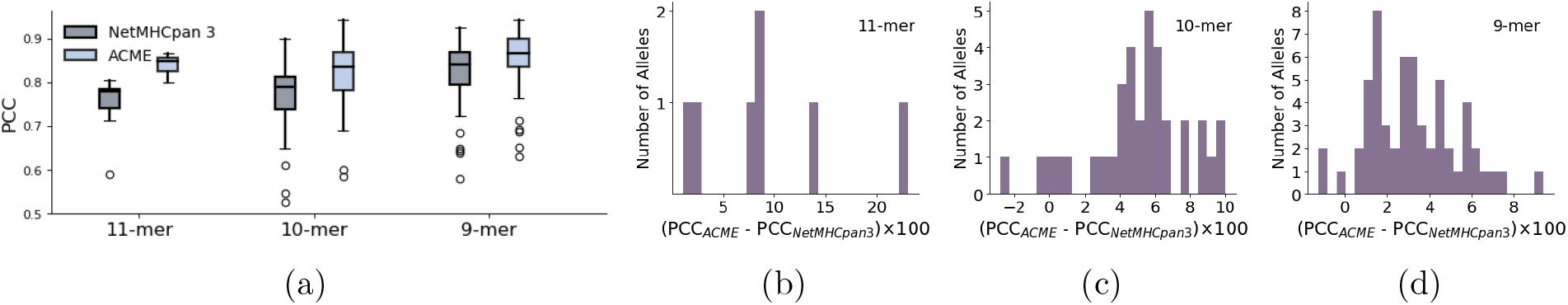
Performance comparison between our model and the state-of-the-art model, NetMHCpan 3.0. a Performances of the two models on peptides of different lengths. In the box plot, each data point represents the Pearson Correlation Coefficient (PCC) value for a specific MHC allele. The prediction results of NetMHCpan were obtained from [8]. b-d Performance improvement of ACME over NetMHCpan 3.0 for 11-mer, 10-mer and 9-mer peptides, respectively. The horizontal axis stands for the improvement (percent), while the vertical axis represents the number of alleles with such improvements.

Note that in the above comparison, ACME was trained mainly on human HLA-peptide binding data while the training data of NetMHCpan 3.0 included additional binding data from other species (which consisted of only 18% of their training data). To show that the improvement of our model was not due to the difference in training data used, we also reimplemented a model with the same model structure and encoding strategy as in NetMHCpan 3.0 [8] (since the source code of NetMHCpan 3.0 was not available), and trained it using the exact same human HLA-peptide binding data as in ACME. The additional comparison based on a five-fold cross-validation also confirmed the significant performance improvement achieved by ACME (Supplementary Table 1).

In addition, we also compared our model to the recently proposed deep learning model, ConvMHC [14], for pan-specific peptide-MHC binding prediction. Note that ConvMHC can only be applied to 9-mers, while our model can also make accurate predictions for peptides with various lengths because our encoding strategy can effectively incorporate critical peptide features that are usually at the two ends of a peptide and is thus insensitive to the variation of peptide length. Moreover, even on 9-mers, ACME still outperformed ConvMHC by 1.2 percent in terms of accuracy. Overall, the above results indicated that ACME can provide significantly more accurate predictions of peptide-MHC binding than other state-of-the-art methods.

### 3.2 Testing the generalizability of ACME

We further conducted two additional tests to demonstrate the generalizability of ACME (Section 2.5). We first tested ACME on 30 other independent datasets, i.e., the most recent IEDB benchmark datasets and compared its prediction performance to those of previous methods, whose Spearman Rank Correlation Coefficient (SRCC) values had been reported previously in [26]. Our comparison showed that the SRCCs of other methods were roughly on the same level, while ACME achieved significantly better performance, with around 5 percent improvement in terms of SRCC (Figure 4a and Supplementary Table 2). These results indicated that our model can be generalized well on other independent data.

**Figure 4:**
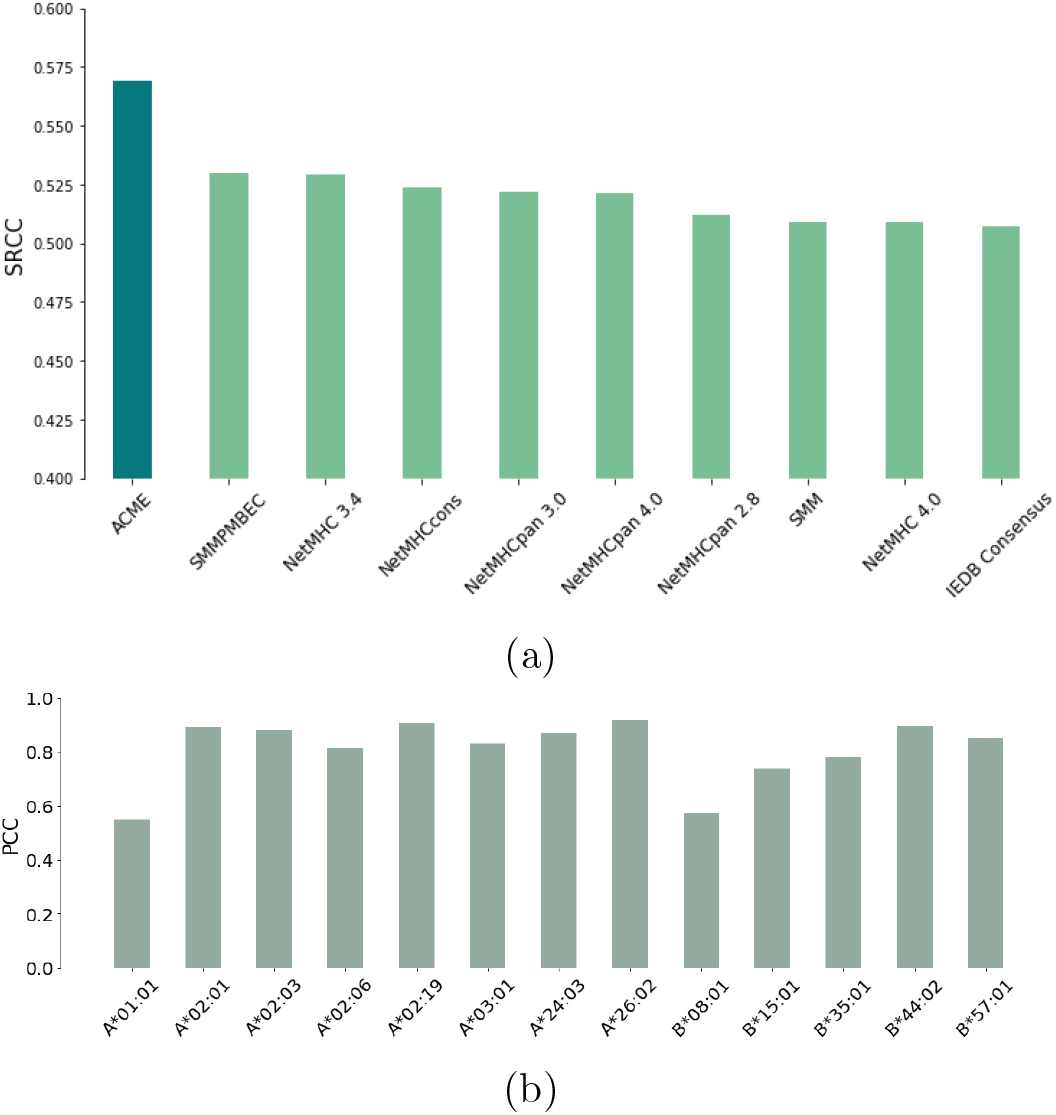
Testing the generalizability of ACME. a The performance of different prediction algorithms on the most recent IEDB benchmark datasets, measured in terms of the Spearman Rank Correlation Coefficient (SRCC). The SRCC scores of other prediction methods were obtained from [26]. Each bar represents the averaged performance of a specific prediction algorithm over all the datasets tested. b The prediction results of the novel alleles that were not seen in the training data. The prediction performance was evaluated in terms of the Pearson Correlation Coefficient (PCC)

Next, we examined whether ACME can make accurate binding predictions for those novel MHC alleles without any training data. Here we selected both common and rare MHC alleles as representatives for testing. As shown in Figure 4b, ACME can still achieve good prediction performance that was close to those alleles that had available training data, with an average PCC score of 0.81 over all the 13 alleles tested (Supplementary Table 3 and Supplementary Notes Section 1.4). This new result indicated that ACME can fully exploit the knowledge from the common alleles with available training data to make accurate predictions for those novel alleles with few or no training data.

### 3.3 Experimental validation

We selected the representative peptides with the highest and lowest predicted binding affinities and experimentally measured their binding affinities to further validate the predictive ability our algorithm.

As shown in Figure 5a, the expression of HLA-A2 was increased when T2 cells were incubated with the peptides with high predicted binding affinities. More importantly, some of the tested peptides with high predicted affinities even displayed equivalent or higher binding affinities when compared to the positive control peptide (OVAL235). Moreover, expectedly, the peptides with low predicted binding affinities also led to a lower expression of HLA-A2 molecules than those with high predicted affinities (*p* = 1.3 *×* 10^−9^, one-sided Mann-Whitney U test).

**Figure 5:**
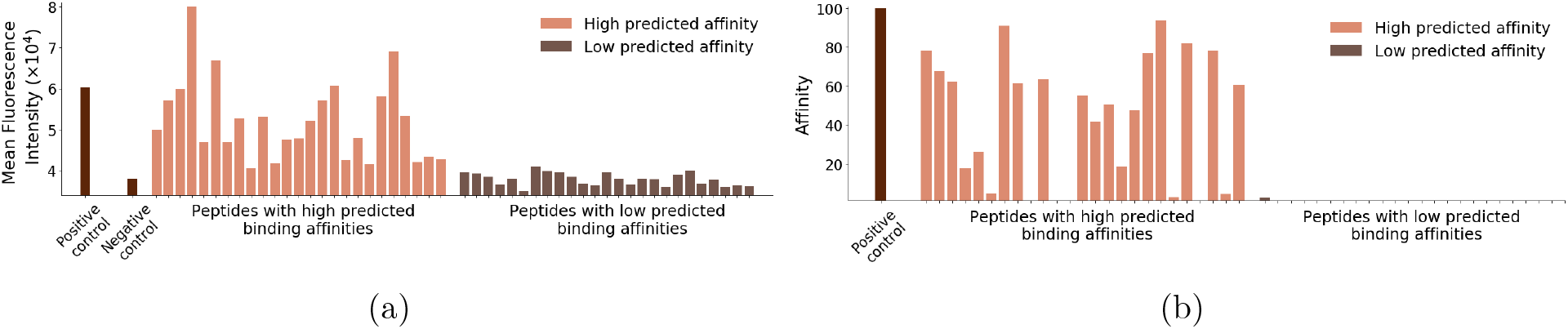
Experimental validation on the peptides with the highest and lowest predicted binding affinities. a The flow cytometry results for HLA-A*02:01, where a higher mean fluorescence intensity (MFI) value indicates a higher affinity. b The ProImmune REVEAL MHC-peptide binding assay results for HLA-B*27:05. Both experiments showed that our binding predictions are generally accurate. Synthesis of one peptide with low predicted binding affinity failed.

We also used the ProImmune REVEAL MHC-peptide binding assay to measure the binding affinities with HLA-B*27:05. The experimental validation results, as shown in Figure 5b, demonstrated that the peptides with high predicted affinities displayed accordingly significantly higher experimentally measured affinities compared to those with low predicted affinities (*p* = 6.1 *×* 10^−9^, one-sided Mann-Whitney U test). Detailed results of the above two experiments can also be found in Supplementary Table 4.

All these experimental validation results further demonstrated that our algorithm can yield accurate predictions and thus provide a practically useful tool to identify neo-antigens in human tumors.

### 3.4 ACME uncovers the underlying rules for MHC-peptide binding

To overcome the black-box nature of deep learning, here we combined our convolutional network with an attention module to enhance the explainability of our model (Section 2.7). To validate the quality of the attention values assigned to each position, we first conducted a masking experiment and confirmed that the positions with higher attention values tend to make a greater contribution to the prediction of MHC-peptide binding in our ACME framework (Supplementary Notes Section 1.5).

As a first attempt, we visualized the motifs for several MHC alleles whose binding patterns had been well characterized previously (Figure 6a). Expectedly, we observed similar preferences for amino acid types at different peptide positions as reported in previous studies [16]. For instance, our model was able to identify the Lys at position 9 as an anchor residue, i.e., the residue making major contributions to binding, for the peptide bound to HLA-A*11:01. Also, for HLA-B*40:01, the previously reported anchor residue Glu at position 2 and Leu at position 9 were also successfully identified by our model as important residues.

**Figure 6:**
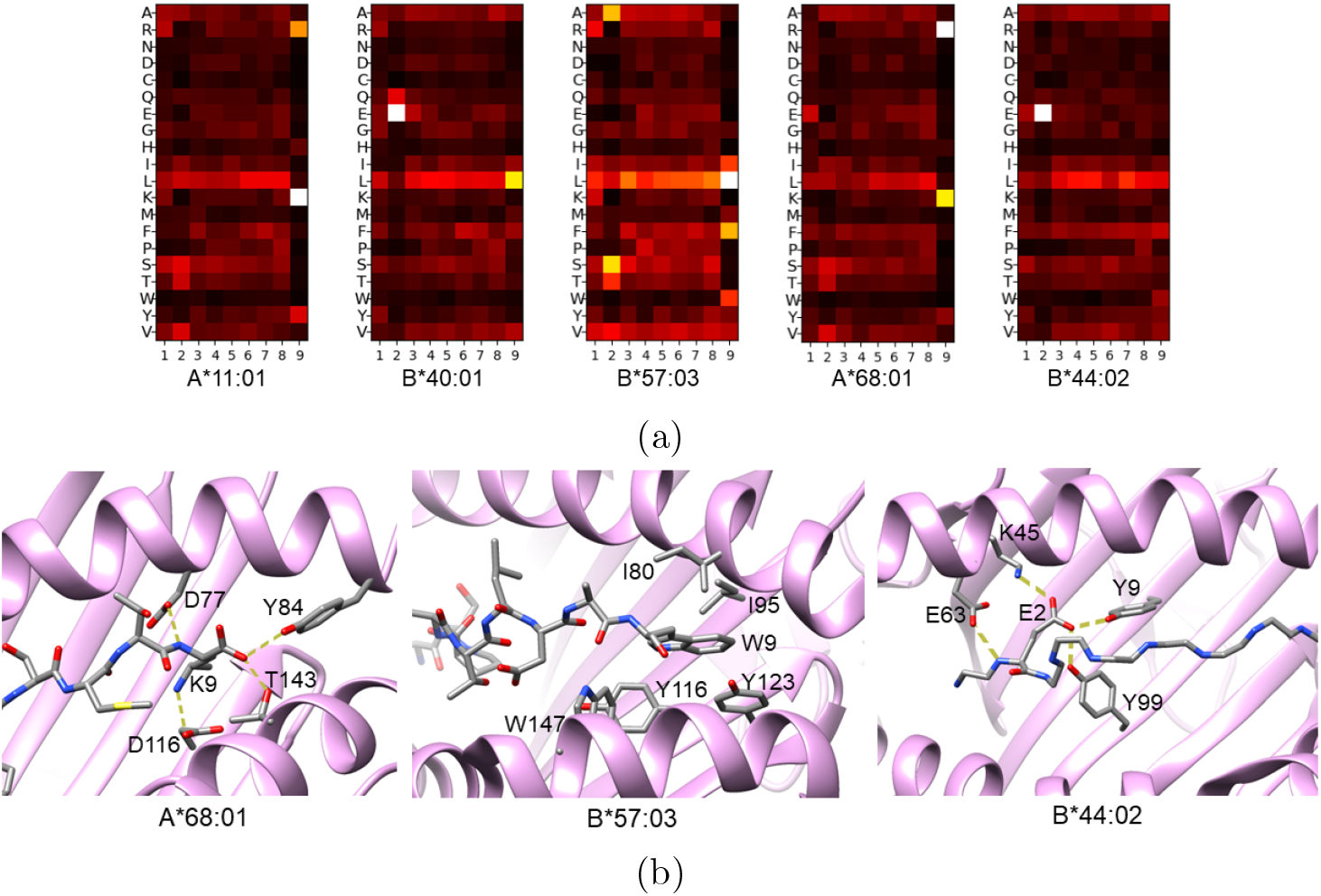
Example of sequence motifs for the peptides bound to different MHC alleles. a The heatmaps visualizing representative motifs for several well-characterized MHC alleles. b Structural basis for the peptide sequence motifs generated by ACME.

In addition, by aligning the motifs discovered by ACME with the previous structural characterizations of MHC-peptide complexes (Figure 6b), we found that our model was also able to capture the local biochemical environment difference caused by the polymorphisms in different MHC molecules, especially at the termini of the peptides. For instance, our analysis demonstrated the Lys or Arg residue at position 9 of a peptide binder significantly contributes to peptide binding to HLA-A*68:01, which can actually be supported by a previously solved high-resolution structure of HLA-A*68:01 complexed with a 9-mer peptide (PDB code 4HWZ) [35], where the C terminus Lys of the peptide is involved in a hydrogen bond network with surrounding residues in the bound MHC molecule. In particular, the *ε*-amino group of this Lys forms a hydrogen bond with Asp116 of the MHC molecule. In contrast, for the HLA-B*57:03 protein whose binding pocket is generally formed by hydrophobic residues, our motif analysis indicated a preference for hydrophobic residues, e.g., Leu, Phe or Trp at the C terminus, which is consistent with the previous structural data (PDB code 2BVP) [36] showing that the C-terminus Trp of the peptide is involved in an aromatic stacking with Tyr123 and makes close contacts with several residues in the MHC molecule, including Trp147, Tyr116, Ile95 and Ile80. In addition, we also found that for our predicted peptides bound to allele HLA-B*44:02, the anchor residue Glu2 near the N terminus is engaged in multiple hydrogen bonds with neighboring MHC residues, and that its *γ*-carboxyl group forms a salt bridge with the *ε*-amino group of Lys45 (PDB code 1M6O) [37]. All these analyses results based on the existing structural data supported the biological relevance of the binding motifs generated by ACME.

Next, we sought to extend this analysis to the MHC alleles whose binding preferences lack experimental characterization. To test the validity of our result, we first confirmed that our model can produce verified motifs for several alleles that were not included in the training data (Supplementary Figure 8a). We then used ACME to generate motifs for a large number of alleles, including those not previously characterized. In particular, we found that positions 2 and 9 in the peptide play the most vital roles in MHC-peptide binding, which was consistent with the findings of previous studies [38]. Our analyses showed that the aliphatic amino acid Leu and Val, the aromatic amino acid Phe and Tyr and the basic amino acid Lys and Arg are among the most common anchor residues. In addition, we found that different alleles may employ different anchor strategy to facilitate specific peptide binding. Specifically, some alleles, e.g., HLA-A*02:19 and HLA-A*24:02, have anchor residues at both ends, while other alleles, e.g., HLA-A*03:02, HLA-A*11:01 and HLAA*11:02, are mainly anchored at one end. We also found that the residues in the middle can also make a significant contribution to binding in some alleles, as can be seen in HLA-B*15:42.

In addition to the discovery of the residues that strengthen binding, we are also interested in identifying the residues that may impair MHC-peptide binding. Therefore, in contrast to the above analyses which focused on high-affinity binding peptides, we also generated a set of non-binder motifs using the peptides with the lowest predicted affinities (Supplementary Figure 8b). These non-binder motifs clearly show foci of attention at several positions, including Ser at position 9 in HLA-A*02:01 and Lys at position 7 in HLA-B*35:01, indicating that most likely these residues are detrimental to binding. To test whether or not these residues can actually impair binding, we conducted an additional test in which we replaced the original residues of the peptides with random amino acids or the above found anti-binding amino acids, respectively. It turned out that replacing the corresponding residues with these anti-binding amino acids indeed resulted in significantly lower predicted binding affinities compared to replacement by random amino acids (Supplementary Figure 7, *p* = 4.3 *×* 10^−185^ for HLA-A*02:01 and *p* = 2.9 *×* 10^−98^ for HLA-B*35:01, one-sided Mann-Whitney U test). This additional test result can thus support our discovery of the non-binder motifs.

## 4 Conclusions

In this study, we have developed a new pan-specific algorithm, called ACME, for peptide-MHC class I binding prediction. The algorithm uses convolutional neural networks to integrate the features extracted from both peptide and the MHC sequences and also has a built-in attention module to help extract interpretable binding patterns. Comprehensive tests on available benchmark datasets have demonstrated the superior prediction performance of ACME over other state-of-theart methods. In addition, we have shown that ACME can be used to accurately generate the binding motifs for different MHC alleles, which can be further used to investigate the patterns of peptide-MHC class I binding. In the future, we will try to incorporate more information such as peptide transportation and TCR recognition to improve the prediction of antigen peptides. We expect that incorporating information about these biological processes into our prediction model will further extend the application potentials of our model in cancer immunotherapies.

